# TMTPro Complementary Ion Quantification Increases Plexing and Sensitivity for Accurate Multiplexed Proteomics at the MS2 Level

**DOI:** 10.1101/2020.10.13.338244

**Authors:** Alex Johnson, Michael Stadlmeier, Martin Wühr

**Affiliations:** Department of Molecular Biology, Princeton University, Princeton, NJ, USA; Lewis-Sigler Institute for Integrative Genomics, Princeton University, Princeton, NJ, USA; Department of Chemical and Biological Engineering, Princeton University, Princeton, NJ, USA

## Abstract

Multiplexed proteomics is a powerful tool to assay cell states in health and disease, but accurate quantification of relative protein changes is impaired by interference from co-isolated peptides. Interference can be reduced by using MS3-based quantification, but this reduces sensitivity and requires specialized instrumentation. An alternative approach is quantification by complementary ions, which allows accurate and precise multiplexed quantification at the MS2 level and is compatible with the most widely distributed instruments. However, complementary ions of the popular TMT tag form inefficiently and multiplexing is limited to five channels. Here, we evaluate and optimize complementary ion quantification for the recently released TMTPro tag, which increases plexing capacity to eight channels (TMTProC). We find that the beneficial fragmentation properties of TMTPro increase quantification signal five-fold compared to TMT. This increased sensitivity results in ~65% more proteins quantified compared to TMTPro-MS3 and even slightly outperforms TMTPro-MS2. Furthermore, TMTProC quantification is more accurate than TMTPro-MS2 and even superior to TMTPro-MS3. To demonstrate the power of TMTProC, we analyzed a human and yeast interference sample and were able to quantify 13,290 proteins in 24 fractions. Thus, TMTProC advances multiplexed proteomics data quality and widens access to accurate multiplexed proteomics beyond laboratories with MS3-capable instrumentation.

## Introduction

Quantitative multiplexed proteomics has become a powerful tool to analyze the proteome across various conditions. Multiplexed proteomics relies on isobaric tags, which are indistinguishable in the MS1 spectrum, but differentiate multiple conditions upon fragmentation in the MS2 or MS3 spectrum (Fig. 1A).^1–3^ Multiplexing is especially attractive because of the increased sample throughput - with current commercial isobaric tags up to 16 conditions can be compared in a single experiment, thus helping to save expensive mass spectrometer instrument time.^2,4^ The different quantification channels are encoded by the distribution of heavy isotopes between the reporter and balancer region of the reagents. Because the overall number of heavy isotopes is kept constant between the different tags, they add the same total mass to the peptides, and hence are isobaric. However, during gas-phase fragmentation, the reporter and balancer group are separated, revealing differences in the masses of each region. This information is used for determining which condition the ions stem from and permits relative quantification of peptides.

**Figure 1:**
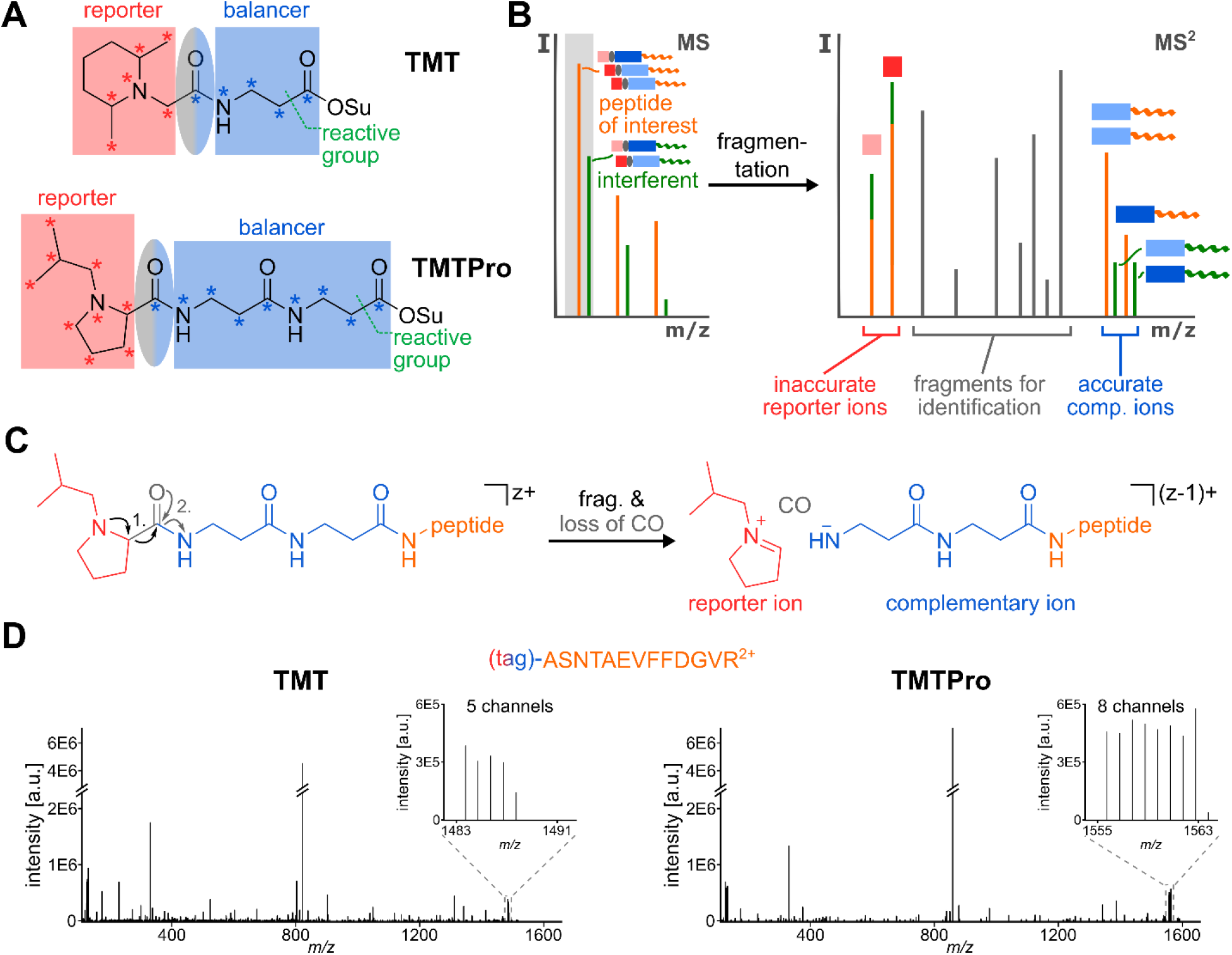
Complementary Ion Quantification with TMT and TMTPro. **A)** TMT- and TMTPro-tags are comprised of a reporter region (red), a balancer region (blue) and an amine-reactive NHS-ester moiety. The carboxyl group lost as CO during fragmentation is part of the balancer and highlighted in an ellipse (grey-blue). While the original TMT-tag can incorporate a maximum of five heavy isotope labels (asterisks) in its balancer group, this number is increased to nine in the TMTPro balancer region. To make use of the increased plexing potential, the reporter group in TMTPro utilizes an iso-butylpyrrolidine-moiety, which can incorporate nine heavy isotope labels. **B)** When analyzing complex samples via shotgun proteomics, in addition to the peptide of interest (orange), other peptides with similar m/z-ratio (interferents, green) will be co-isolated (grey box). If MS2 reporter ions are used for quantification, the interfering peptides lead to a distortion of the measured ratios, as the source of reporter ions (red squares) cannot be distinguished. However, because the masses of complementary ions are peptide-dependent and include the heavy isotope labels of the balancer-region (blue rectangles), they can be used for interference-free, accurate MS2 quantification. **C)** During fragmentation of a TMTPro-modified peptide, the positively charged reporter ion is separated from the ion and a neutral CO-molecule is lost. This leads to an ion where the balancer part is still attached to the peptide. Because the balancer region encodes the complementary heavy isotope labels of the reporter ion, the balancer-peptide conjugate is called the TMTProC or complementary ion. In this process, the charge state of the precursor ion is reduced by one. **D)** Example MS2 spectra of the peptide ASNTAEVFFDGVR^2+^, labeled with TMT (left) or with TMTPro (right) and fragmented with HCD. Both MS2 spectra exhibit a very similar total ion count. However, as the inserts show, TMTPro-labeled peptides generate eight instead of five complementary ions with higher absolute intensity.

An inherent advantage of multiplexed proteomics is that the samples are co-analyzed, avoiding problems with missing values that are common in label-free experiments. The co-analysis of all samples in the same MS experiment enables exquisite measurement reproducibility and precision. Because isobaric tags are attached after sample lysis, data collection is compatible with essentially any protein sample, avoiding the need for isotopic labeling in living systems, which is common in methods that depend on heavy isotopes (e.g. SILAC).^5^ These advantages have led to the ever increasing popularity of multiplexed proteomics, leading to a wide variety of findings in breast cancer treatment,^6^ lung cancer metastasis,^7^ as well as fundamental research into translation regulation,^8^ among many others.

A major challenge inherent to multiplexed proteomics quantification is measurement accuracy. In its simple implementation, co-eluting peptides with similar mass to charge ratios are co-isolated and co-fragmented with the peptide of interest. The resulting quantification is typically significantly distorted (Fig. 1B).^1,3,9,10^ The most widely used method to overcome ratio distortion uses an additional MS3 scan.^1,11^ In these methods, commercialized on Thermo Fisher tribrids,^12^ several b- and y-ions from the MS2 scan are simultaneously co-isolated and fragmented in an ion-trap (multi-notch or SPS-MS3). The extra gas-phase purification step leads to a significant decrease of ratio distortion. However, this advantage comes at the cost of decreased sensitivity and the need for highly specialized instrumentation with MS3 capabilities, which are just a small fraction of proteomics capable instrumentation currently in use.

An alternative method for accurate multiplexed proteomics is to make use of the complementary ions.^13,14^ This method was designed as a quantification strategy that does not require higher order scans and can be employed on a wide variety of instruments. When peptides are fragmented in the MS2 spectra, the loss of the isobaric reporter ion leaves the balancer region with a complementary isotope distribution behind. These balancer-peptide-conjugates, the so-called complementary ions, are peptide-dependent and typically have a slightly different mass than co-isolated peptides (Fig. 1A, B). Therefore, using the complementary ions for quantification drastically reduces ratio distortion effects compared to both MS2 reporter ion quantification as well as multi-notch MS3 approaches. Further improvements to the method, including a narrower isolation window and modelling of the isolation window shape in the deconvolution algorithm, further improved measurement precision.^14^ This approach, called TMTc+ when used with the TMT isobaric tag, has been successfully applied to multiple biological research studies.^15–18^

Despite its attractiveness, remaining challenges hinder widespread application of TMTc+. First, the plexing capacity of TMTc+ is limited to five channels because the small mass differences between ^13^C and ^15^N cannot be resolved in the high *m/z*-regime of the complementary ions. In addition, the loss of CO during TMT-fragmentation reduces the number of heavy isotopes available for encoding quantification channels by one. Furthermore, commercial isobaric tags were not designed for this approach and complementary ion formation is comparatively poor. High energies are necessary to separate the reporter from the balancer region, but the peptide backbone is also amenable to breaking at these levels, leading to reduced complementary ion intensity.

Easy to cleave sulfoxide tags, like the SO-tag and EASI-tag, were designed to improve complementary ion formation efficiency.^19,20^ However, fragmentation with these tags happens too readily, typically leading to additional fragmentations and MS2 spectra of very high complexity, which hinders identification and leads to low identification success-rates and distributing much of the signal away from the intact peptide complementary ions. Though we believe these processes generate complement fragment b- and y-ions that might be very attractive for multiplexed Data-Independent Acquisition or targeted Data-Dependent Acquisition approaches,^3^ they provide a severe hindrance for shotgun-proteomics.

Recently Thermo released a new isobaric tag, named TMTPro.^2^ This tag was primarily designed for Thermo’s MS3 methods and is commercially available as encoding up to 16 different conditions in a single experiment. We noticed that this proline-based tag breaks easier than the previous TMT-tag while not having a detrimental effect on identification rates. We reasoned that this tag could be well-suited for the complement reporter ion approach. Here, we optimize data acquisition strategies for quantification of complement reporter ions with TMTPro in a method termed TMTProC. We find that TMTPro significantly improves the efficiency of complementary ion formation compared to TMT and increases plexing from five to eight with one Dalton separation. TMTProC also reduces ratio compression more than MS3 methods while maintaining sensitivity levels equivalent to TMTPro-MS2.

## Methods

### Proteomics sample preparation

Samples were mostly prepared as previously described.^14,21,22^ Human peptides from HeLa cell lysate and yeast peptides from *Saccharomyces cerevisiae* lysate were both used for method optimization. For description of lysis conditions and further information, please reference the supporting information.

Briefly, lysates were reduced by 5 mM DTT (20 min, 60 °C), alkylated with 20 mM NEM (20 min, RT), and quenched with 10 mM DTT (10 min, RT). Proteins were purified by methanol-chloroform precipitation^23^ and afterwards resuspended in 10 mM EPPS pH 8.0 with 6 M guanidine hydrochloride (GuHCl). They were then diluted to 2 M GuHCl with 10 mM EPPS pH 8.0 and digested with 10 ng/μL LysC (Wako) at room temperature overnight. Samples were further diluted to 0.5 M GuHCl with 10 mM EPPS pH 8.0 and digested with an additional 10 ng/μL LysC and 20 ng/μL sequencing grade Trypsin (Promega) at 37 °C for 16 hours.

Samples were then vacuum dried and re-suspended in 200 mM EPPS at pH 8.0. TMTPro tags were mixed at the appropriate ratios prior to labelling peptides. The TMTPro mixture was added at a molar tag:peptide ratio of 5:1 and allowed to react for 2 hours at room temperature. The reaction was quenched with 1% hydroxylamine (30 min, RT). Peptides were then acidified to pH~2 with phosphoric acid.

Unfractionated samples were centrifuged at 24k rcf for 10 minutes at 4 °C before desalting *via* SepPak-cartridges (Waters). Samples were vacuum-dried and re-suspended in 1% formic acid before mass spectrometer analysis. For pre-fractionation, samples were spun at 50krcf for 1 hour at 4 °C and separated by reverse-phase HPLC at high pH. For samples used to evaluate ratio accuracy and the extent of peptide interference, human and yeast peptides were mixed immediately prior to mass spectrometry analysis.

### LC-MS experiments

Samples were analyzed on an EASY-nLC 1200 (Thermo Fisher Scientific) HPLC coupled to an Orbitrap Fusion Lumos mass spectrometer (Thermo Fisher Scientific), with Tune version 3.3. Peptides were separated on an Aurora Series emitter column (25 cm x 75 μm ID, 1.6 μm C18) (ionopticks, Australia), held at 60 °C during separation by an in-house built column oven, over 120 min for unfractionated and 90 min for fractionated samples, applying nonlinear acetonitrile-gradients at a constant flow rate of 350 nL/min. Samples were analyzed with either an MS2-CID-method for TMTProC or with conventional MS2-HCD- and SPS-MS3-methods adjusted for optimal performance for reporter ion-based quantification.

To investigate the effect on ratio distortion when utilizing field asymmetric ion mobility spectrometry, a FAIMS Pro device (Thermo Fisher Scientific) was used in some experiments.^24^ For detailed information about the gradients and methods utilized, as well as data analysis procedures, please see the supporting information.

### Proteomics data availability

The mass spectrometry proteomics data have been deposited to the ProteomeXchange Consortium via the PRIDE^25^ partner repository with the dataset identifier PXD021661 and 10.6019/PXD021661 with username and password vWrS8ZgP.

## Results and Discussion

### TMTPro increases plexing for complementary reporter ion quantification

A major shortcoming of the previous complementary reporter ion approach (TMTc+) was the limit of five encoded channels.^13,14^ Thermo Fisher recently released an isobaric tag (TMTPro) with a new proline-based reporter group and a longer balancer region that can accommodate up to nine heavy isotopes, compared to five heavy isotopes accessible in TMT (Fig. 1A).^2^ The current set of commercially available TMTPro tags increase plexing for the complement reporter ion approach from five (TMT) to eight (Fig. 1D, Sup. Fig. S1A). A structure without heavy isotopes in the balancer region could increase plexing capacity to nine while maintaining the same total number of heavy isotopes (Sup. Fig. S1B).

The slight mass difference between isotopomeric structures incorporating ^13^C vs ^15^N allows up to 16 distinguishable channels with reporter ion quantification in high-resolution mass analyzers (50k resolution at *m/z* 200).^2^ However, at the higher mass range of complementary reporter ions, these small mass differences cannot be resolved while maintaining high acquisition speed. Super-resolution acquisition is a very active area of research and once a resolution of ~60k at the high *m/z*-ranges of complementary ions becomes available,^26,27^ twelve channels could be distinguished with the current set of TMTPro tags (see Sup. Fig. S1A). Theoretically, isotopomeric structures with a total of seven heavy isotopes could encode up to 21 different conditions using TMTProC. The higher plexing capacity of TMTPro for complementary ions using fewer heavy isotopes, compared to low *m/z* reporter ion quantification, is due to the two nitrogen atoms in the balancer region, while there is only a single nitrogen atom in the reporter region. A structure with the same number of total heavy isotopes in the complementary ion could therefore be split into three independent channels.

### TMTProC generates highly accurate quantitative proteomics data

To evaluate the accuracy of measured ratios and distortion due to interfering peptides, we prepared a sample of mixed HeLa lysate and *Saccharomyces cerevisiae* (yeast) lysate (Fig. 2A).^1^ HeLa peptides were labelled with TMTPro in ratios of 1:1 in all eight channels, while yeast peptides were labelled in ratios of 0:1:5:10:10:5:1:0. The two lysates were combined after labelling with ten parts HeLa for every one part yeast peptides. This sample simulates the quantification of lower abundant peptides that change concentration between conditions (yeast) in a background of highly abundant peptides that don’t change concentration (HeLa). When isolating yeast peptides, co-isolation of HeLa peptides will tend to bias the measured channel ratios towards 1, making quantification less accurate. The mixed HeLa-yeast sample was analyzed with a 90-minute run using three different quantification methods: TMTPro-MS2, TMTPro-MS3, and TMTProC. Isotopic impurities in the reporter and complementary regions of each tag were measured using heavy-labelled arginine reacted with TMTPro (see Supporting Information). For accurate determination of the measured ratios, we adapted the TMTc+ deconvolution algorithm for TMTPro.^14^ The source code is available via github (https://github.com/wuhrlab/TMTProC). In addition, we assessed the effectiveness of a High-Field Asymmetric Waveform Ion Mobility Spectrometry (FAIMS) device to reduce the effect of ratio distortion using each method.^24^

**Figure 2.**
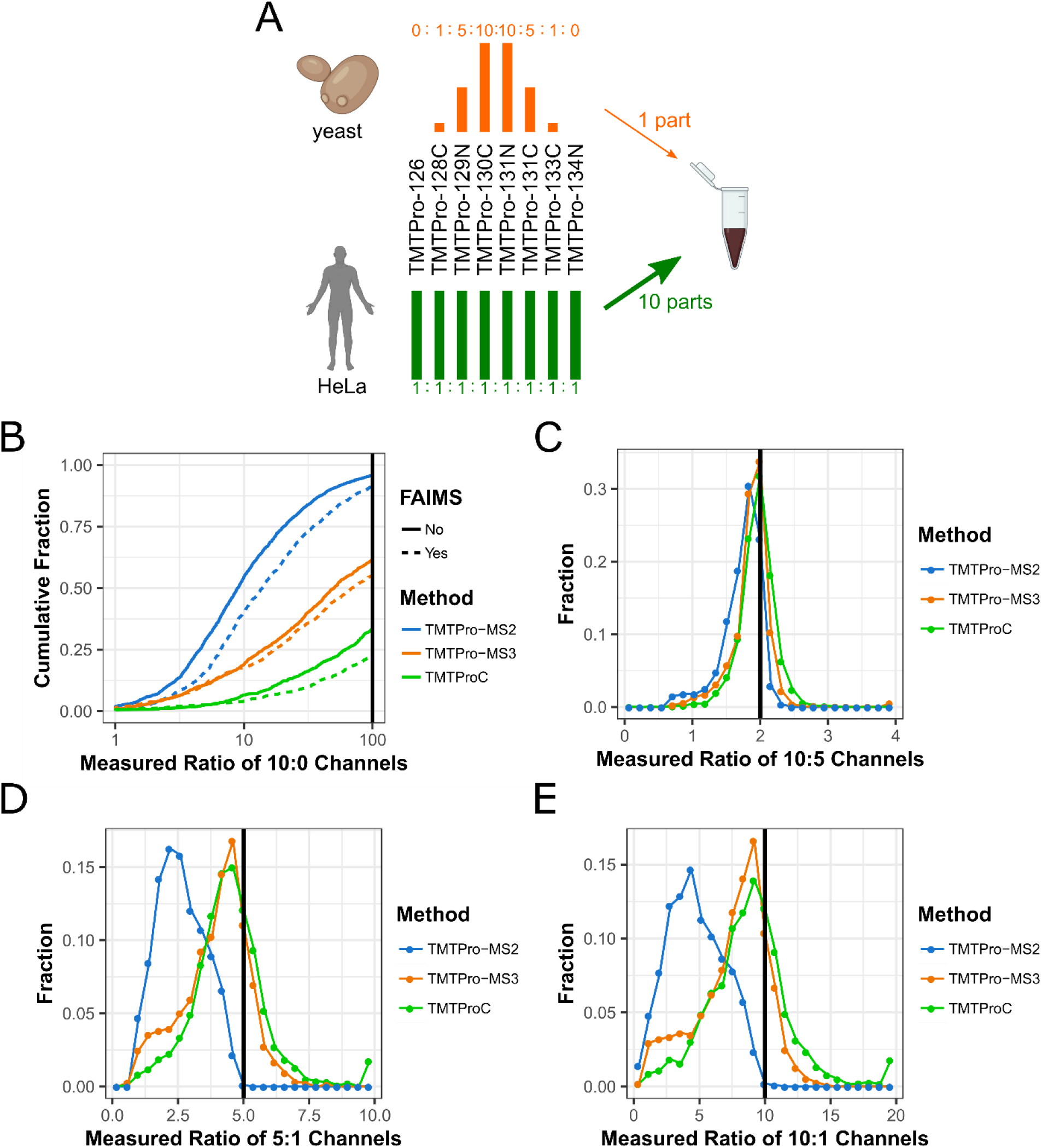
Evaluating ratio distortion for TMTProC and alternative multiplexed quantification strategies. **A)** Yeast lysate labelled with TMTPro in ratios of 0:1:5:10:10:5:1:0 was mixed with HeLa lysate labelled with TMTPro in ratios of 1:1:1:1:1:1:1:1 at a mixing ratio of 1 (yeast):10 (HeLa). **B)** The cumulative distribution function of measured ratios for each peptide of the two 10:0 channel pairs was calculated using three different quantification methods, each with and without ion-mobility prefractionation (FAIMS). Peptides for which the sum of the quantifiable ions signal to noise ratio (S:N) was less than 500 were removed for all methods. Other signal to noise cutoffs can be found in Sup Fig. S5. Measured ratios greater than 100 were set to 100. TMTPro-MS3 significantly reduces interference compared to TMTPro-MS2. TMTProC reduces interference even further compared to TMTPro-MS3. **C-E)** Histograms showing the measured ratios of the 10:5 (C), 5:1 (D), and 10:1 (E) channels for yeast peptides with various quantification strategies. Peptides with a sum S:N of less than 200 were removed from this analysis. Measured ratios outside the histogram were set to the closest ratio shown. MS2 quantification of the 10:1 and 5:1 channels is plagued by interference, which MS3 and TMTProC significantly reduce. TMTProC outperforms TMTPro-MS3, since fewer peptides are significantly distorted (shoulders on the left side of the 5:1 and 10:1 histogram).

In the analyzed sample, ratio distortion from interfering HeLa peptides can be assessed by calculating the ratio of the signal to Fourier transform noise (S:N) of the 10 and 0 channels on each side of the reporter or complementary ion envelope for each yeast peptide. The lower the measured ratios, the more co-eluted HeLa peptides are interfering, while a ratio approaching infinity would be the expected outcome without interference. Peptides with less than 500 total S:N in the complementary or reporter envelope were removed, regardless of the quantification method, so that a S:N ratio of 100 could theoretically be detected from the 10:0 channel. The cutoff ensures that peaks at the noise level are not misinterpreted as a measured infinite ratio due to low signal.

The cumulative distribution function of the relative S:N ratio in the 10:0 channels is presented in Fig. 2B. TMTPro-MS3 (orange) significantly outperforms TMTPro-MS2 (blue), with 38.7% of peptides having a measured ratio greater than 100, as compared to 4.2% for TMTPro-MS2. Decreasing spectra complexity by utilizing a FAIMS device moderately reduced interference for TMTPro-MS2, but not to the level of TMTPro-MS3. The highest reduction of interference is observed with TMTProC, yielding even less interference than TMTPro-MS3, both for runs with and without a FAIMS device. Using TMTProC (green), more than 66% of peptides had a measured ratio greater than 100, which agrees with the results using TMTc+.^14^ FAIMS consistently reduced ratio distortion for all three quantification methods, although the method used had a stronger effect in all cases.

We also evaluated the effect of ratio distortion on the accuracy and precision of the remaining channel ratios by determining the measured ratios for the 10:5 channels, the 5:1 channels, and the 10:1 channels (Fig. 2C-E). TMTPro-MS3 and TMTProC were able to reproduce the expected 10:5 ratio of the innermost channels with only small variations between the methods (Fig. 2C). Even TMTPro-MS2 performed reasonably well for this ratio, but ratio distortion is clearly observable.

Ratio distortion significantly impacts the measured ratio of the 5:1 and 10:1 ratios using TMTPro-MS2. The median ratios for this method were 2.6 and 4.6, respectively (Fig. 2D, E). The use of FAIMS slightly improved the median measured ratios of the channels to 3.1 and 5.9, respectively, for MS2 reporter ion quantification (Sup. Fig. 4), again suggesting that the quantification method has a much stronger effect on quantification accuracy than utilizing a FAIMS device.

TMTPro-MS3 and TMTProC reduce interference significantly for both the measured 5:1 and 10:1 ratios. We observe that the mode for TMTPro-MS3 and TMTProC is slightly distorted compared to the expected ratio (Fig. 2D, E). This could be due to interference or slight errors arising during data normalization. The two methods differ moderately in the tails of their distributions. With these ratios, TMTProC found no signal in the lower abundant channel for ~2% peptides, likely due to interference from peptides with slightly different complementary masses than the peptide of interest, thus shifting the true complementary peak. This effect was negligible for the 10:5 ratio. On the left tail, TMTPro-MS3 performs significantly worse than TMTProC, with 18% of peptides having a ratio less than 2.5:1 for the 5:1 channels (Fig. 2D) and 15% of peptides having a ratio less than 5:1 in the 10:1 channels (Fig. 2E). With TMTProC, the amount of these peptides was nearly cut in half, with 7% of 5:1 channels having a ratio less than 2.5:1 (Fig. 2D) and 10% having a ratio less than 5:1 for the 10:1 channels (Fig 2E). Overall, these results show that TMTProC reduces interference from contaminating peptides as well as or better than MS3 quantification, resulting in highly accurate quantification even in highly complex samples.

### TMTPro efficiently forms complementary ions

So far, we have shown that TMTPro increases plexing and maintains superior measurement accuracy for the complementary reporter ion strategy. Next, we evaluated how efficiently TMTPro forms complementary ions, which is a major shortcoming of TMT for this strategy.^13^ For TMT, this is particularly challenging for higher charge states and for ions that contain highly mobile protons, i.e. more charges than are localized on arginine, lysine, and histidine. In previous studies we selected only 2+ ions for complementary reporter ion quantification with TMT,^13-16,21^ due to inefficient formation for ions with z = 3. We wondered if TMTPro would break more easily and increase the flux into complementary ions which would further increase the method’s sensitivity.

First, we optimized various fragmentation methods and energies for TMTPro complementary reporter ion formation. To do so, we labeled HeLa peptides with TMTPro without heavy isotopes (TMTPro0). For each quantified peptide we calculated the ratio of the ion flux in the precursor peak of the MS1 spectrum and the ion flux of complimentary ions in the MS2 spectrum (see Supporting Information).

Using beam-type HCD fragmentation^28^, we find that the optimal normalized collision energy for complementary ion generation is around 29% for 2+ ions, while 3+ ions had a slightly lower optimal energy of 27%. Both optimal energies were lower than those used for TMT owing to the relatively facile fragmentation of the TMTPro reagent. Peptides with a 2+ charge transmitted on average more than 11% of precursor ion flux in the MS1 into the complementary ion peak of the MS2. Peptides with a 3+ charge formed complementary ions at around a 6% median efficiency, a nearly 2-fold reduction. Both of these efficiencies exceed that measured for peptides with a 2+ charge state tagged with TMT at only 4% median efficiency. Peptides with a charge state of 4+ and 5+ were also considered (data not shown), but efficiencies were too low to warrant further investigation.

We also explored resonance collision induced dissociation (CID)^29^ of TMTPro. With CID, we see a nearly two-fold increase in complementary ions over HCD for both 2+ and 3+ ions (Fig. 3A). There was no relationship between normalized CID collision energy or activation time and complementary ion formation efficiency over the range of 25-31% energy (data not shown). The combination of TMTPro and CID fragmentation increased total complementary ion formation efficiency by 5-fold over TMT fragmented with the optimized method (32% HCD). This major improvement brings TMTProC close to the efficiencies of low *m/z* reporter ion-based quantification methods, significantly increasing the sensitivity of the complementary reporter ion quantification approach.

**Figure 3.**
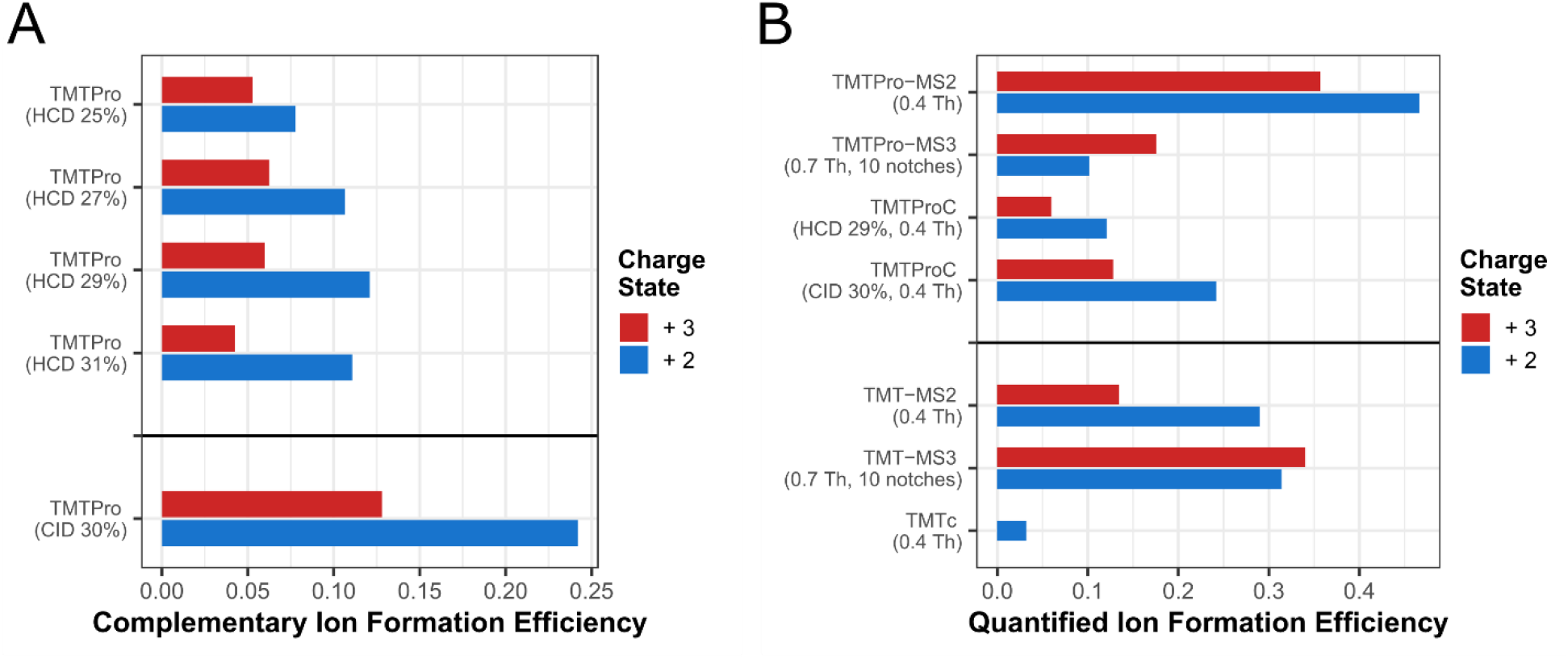
Optimization of TMTProC ion formation and comparison with ion formation efficiencies in alternative multiplexed proteomics methods. **A)** Evaluation of various fragmentation methods for TMTProC ion formation. HeLa peptides were labelled with TMTPro and subjected to various fragmentation modalities. Shown are the median proportions of MS1 ion flux from the precursor that was converted into complementary ions. CID fragmentation produced about two-fold higher signal in the complementary ion cluster than HCD fragmentation. **B)** Comparison of TMTProC ion efficiencies with alternative methods. HeLa peptides labeled with TMTPro or TMT were subjected to various quantification methods. For each, we calculate the median fraction of precursor ion flux that is converted into the relevant ion used for quantification. TMTPro improves complementary ion formation efficiency to the same level as both reporter ion-based quantification methods.

### Efficient TMTProC ion formation results in high sensitivity for complex quantitative proteomics studies

A significant advantage of TMTProC over TMTPro-MS3 is the increased depth of proteome coverage. Using a sample of TMTPro0-tagged HeLa peptides, we quantified 3,824 proteins in a single 2-hour run at 1% protein false discovery rate (FDR) using CID fragmentation. The equivalent run with TMTPro-MS2 quantified 3,361 proteins, while TMTPro-MS3 only quantified 2,295 proteins (Fig. 4A). TMTProC with HCD fragmentation did not perform as well as CID fragmentation but was still very competitive with TMTPro-MS2 (3,000 quantified proteins). Similarly, TMTProC quantifies significantly more peptides than TMTPro-MS3 (18,145 vs. 11,482), and a similar number of peptides as TMTPro-MS2 (16,796) (Fig. 4B). The reduction in sensitivity for TMTPro-MS3 is almost entirely due to the higher overhead times of performing an additional MS scan. Previous studies have shown that a real-time search (RTS) algorithm can improve the efficiency of MS3-based methods by obviating the need for MS3 scans where no peptide could be matched to the MS2 spectrum.^30^ The mass spectrometer used in this study is not capable of RTS, so we cannot compare the sensitivity of this method to TMTProC. We expected that TMTPro-MS2 would be more sensitive than TMTProC because reporter ions are formed at higher efficiency, thereby lowering injection times and increasing duty cycles. However, this effect is offset by the need to filter peptides from the TMTPro-MS2 run where the isolation specificity is less than 75% to diminish interference effects, as peptides with low isolation specificity are subject to more interference than those with higher isolation specificity (Fig. 4C).^11^ Filtering TMTProC data for this criterion is unnecessary because interfering peptides are inherently excluded from quantification.

**Figure 4.**
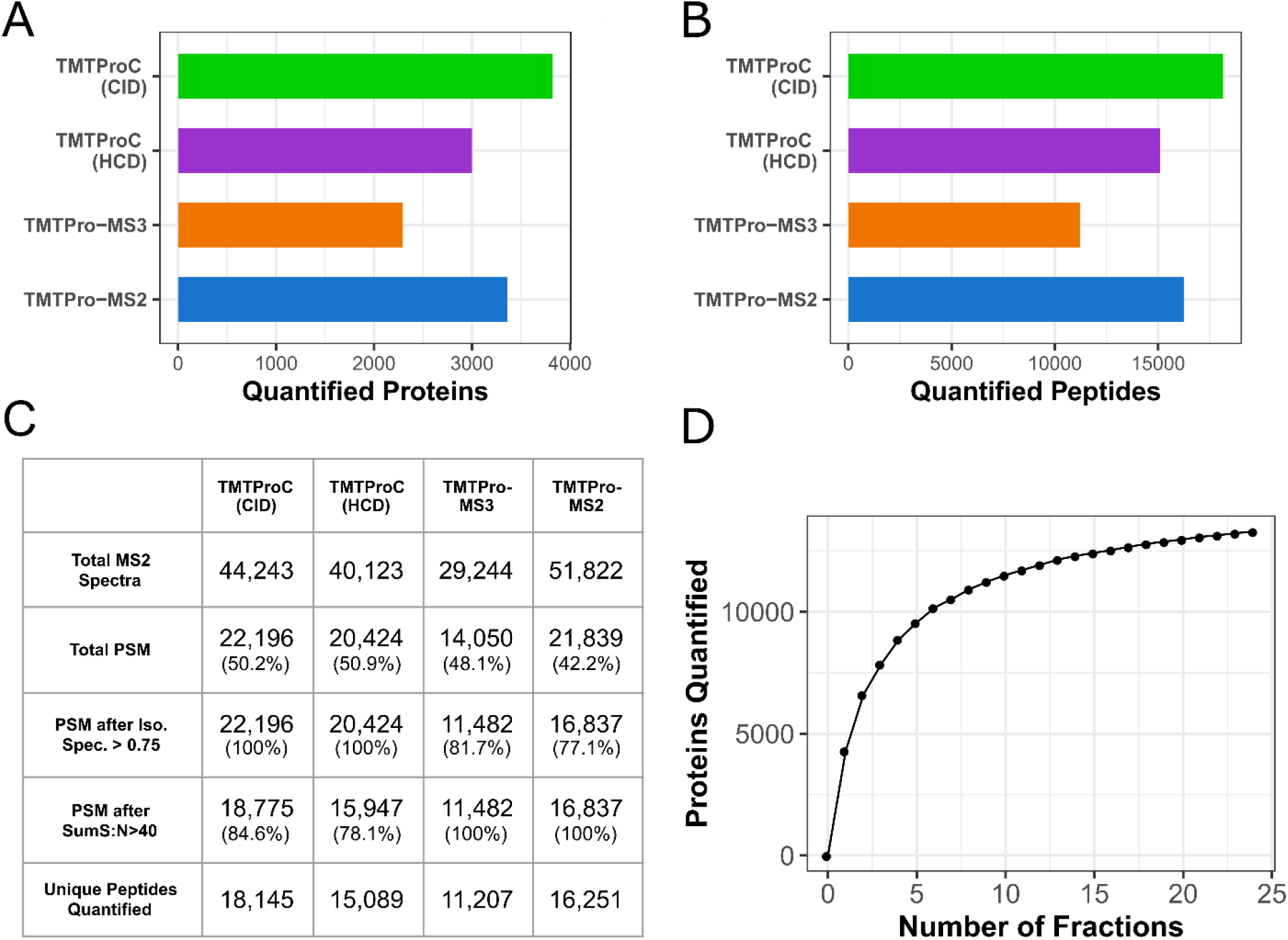
Evaluating TMTProC sensitivity for the analysis of a sample proteome. **A, B)** Number of quantified HeLa proteins (A) and peptides (B) in a single 120 min unfractionated analysis of a TMTPro0-tagged HeLa lysate sample. TMTProC using CID fragmentation is more sensitive than either TMTPro-MS3 or TMTPro-MS2 in the number of quantified peptides and proteins. The sensitivity of TMTProC using HCD fragmentation significantly outperforms TMTPro-MS3 and is competitive with TMTPro-MS2 at the peptide level. **C)** Peptide spectral matches (PSM), quantifications and effects of filter criteria for the analyses in A) and B). Percentages in parentheses in each cell represent the proportion of PSM that passed that filter. With TMTProC, isolation specificity filters do not have to be applied due to higher interference resistance, but stringent Sum S:N thresholds for the ions used for quantification lead to some peptide removal. Still, after filtering, TMTProC outperforms the reporter ion methods in both number of quantified peptides and overall quantification rate. **D)** Number of proteins quantified from a mixed sample of HeLa and yeast peptides shot with TMTProC (S:N>40, 1% Protein FDR, max ppm deviation<10) as a function of the number of fractions used. Only a handful of fractions were necessary to quantify more than 10,000 proteins, and the full 24 fractions led to 13,960 quantified proteins.

We chose to consider a peptide as quantified if its complementary ion envelope signal to FT noise (S:N) summed to at least 40. Recent advancements in integrating ion statistics with peptide concordance allow for peptides with nearly any S:N to improve ratio estimates.^21^ We therefore used 40 S:N as a conservative cutoff for this study. To use this Bayesian inference method (BACIQ), we need to be able to convert signal to noise into pseudo-counts. We have done so for peptides with 2+ and 3+ charge states at various Orbitrap resolutions as shown in Supplemental Figure S3.

To test the limits of TMTProC sensitivity, we prefractionated a 1:1 mixture of human and yeast peptides by mid-pH reverse-phase HPLC into 24 fractions. Each fraction was analyzed with a 90-minute gradient. Across all fractions, we quantify 13,290 proteins at a 1% protein FDR and signal:noise cutoff of 40 (Fig. 4D). Peptides were also removed if the measured *m/z* value of any complementary peak disagreed with their expected value by more than 10 ppm. 4,610 of these quantified proteins were yeast and 8,680 were from HeLa. A similar sample of HeLa peptides, also separated into 24 fractions, which was analyzed using TMTc+, resulted in 8,943 protein quantifications.^14^ A mixed human and yeast sample using the EASI tag quantified 9,760 proteins using four-fold more instrument time and less stringent quantification filtering criteria.^20^ These results demonstrate the significantly improved sensitivity of TMTProC over other complementary ion-based quantification strategies.

## Conclusion

In this paper we have demonstrated that the combination of TMTPro with complementary ion quantification for multiplexed proteomics, termed TMTProC, has superb accuracy, sensitivity, higher plexing capacity, and is subject to relatively little interference from co-eluting peptides. We show that TMTProC maintains the ratio precision of TMTc+, and owing to more facile fragmentation, forms complementary ions for quantification at nearly 5-fold the efficiency of TMT. Resonance CID and beam-type HCD fragmentation are both considered, with CID fragmentation outperforming HCD fragmentation for complementary ion formation. With the increase in complementary ion formation efficiency, we also recommend the isolation of 3+ ions in addition to 2+, a strategy which was not feasible using TMT. Use of a FAIMS device slightly reduced interference from co-eluting peptides in all cases, but the effect of the quantification method was more pronounced. We utilized the optimized method to quantify 13,290 proteins in 24 fractions each analyzed with a 90-minute gradient.

Furthermore, TMTProC can be performed on relatively simple instruments compared to multinotch MS3 methods. The method is compatible with multiple mass spectrometers, e.g. with the QExactive/Exploris platform or QTOF-platforms, encompassing the majority of commercial proteomics instruments. Therefore, TMTProC opens up the possibility of high-quality proteomics measurements to many research laboratories that were previously reliant on inaccurate, interference prone MS2 reporter ion quantification.

## Acknowledgments

We would like to thank Lance Martin (supported by the Princeton Catalysis Initiative). We thank Meera Gupta, Thao Nguyen and Felix Keber for helpful comments on the manuscript. We thank Graeme McAlister for helpful suggestions and discussions. This work was supported by the Eric and Wendy Schmidt Transformative Technology Fund, NIH grant R35GM128813, and the U.S. Department of Energy, Office of Science, Office of Biological and Environmental Research under award number DE-SC0018420.

## Supporting information

### Sample preparation

Samples were essentially prepared as previously described.^14,22^ HeLa S3 cells were grown on 10 cm tissue culture plates to 80% confluency, and yeast cells were grown to an OD of 1.0 in suspension. HeLa cells were pelleted and lysed by sonication in 100 mM HEPES buffer at pH 7.2 with 2% SDS and Roche protease inhibitor. Yeast cells were lysed by cryomilling, and later re-suspended in 50 mM HEPES at pH 7.2 with 4% SDS and 1 mM DTT.

Lysates were diluted to 2 μg/μL with 100 mM HEPES pH 7.2. For reduction of disulfides, DTT (500 mM in water) was added to a final concentration of 5 mM (20 min, 60 °C). Samples were cooled to room temperature and cysteines were alkylated by the addition of N-ethyl maleimide (NEM, 1 M in acetonitrile) to a final concentration of 20 mM and incubation for 20 min at room temperature. 10 mM DTT (500 mM stock, water) was added at room temperature for 10 min to quench any remaining NEM. For protein clean-up, a methanol-chloroform precipitation was performed. The resulting protein pellet was washed with 50/50 methanol/chloroform one additional time and the protein was allowed to air dry.

Protein samples were dissolved in 6 M guanidine hydrochloride, 10 mM EPPS pH 8.5 to ~4.5 μg/μL. Samples were heated at 60 °C for 15 min to help resolubilization and protein content was determined by BCA assay (Pierce BCA Protein Assay Kit, Thermo Scientific).

Next, an appropriate amount of protein (350 - 450 μg) was diluted with 10 mM EPPS pH 8.5 to 2 M Guanidine hydrochloride. Lysates were digested overnight at room temperature with LysC (Wako, 2 μg/μL stock in HPLC water) at a concentration of 10 ng/μL LysC. Samples were further diluted to 0.5 M guanidine hydrochloride with 10 mM EPPS pH 8.5, an additional 10 ng/μL LysC as well as 20 ng/μL of sequencing grade Trypsin (Promega) were added and samples were incubated at 37 °C for 16 hours. Samples were vacuum-dried and then resuspended in 200 mM EPPS, pH 8.0 to a peptide concentration of 1 μg/μL.

To limit experimental noise due to labelling efficiency, TMTPro tags were mixed at the appropriate ratios prior to labelling peptides. 25 μL of the appropriate TMT-reagents (Pierce, 20 μg/μL in dry acetonitrile stored at −80 °C) was added for each 100 μg of peptides, mixed, and incubated at room temperature for 2 hours. The reaction was quenched by addition of 10 μL of 5% hydroxylamine for each 100 μg of peptide (Sigma, 50% in water, HPLC grade, diluted with HPLC water) at room temperature for 15 minutes.

This mixture was acidified to pH < 2 with phosphoric acid (HPLC grade, Sigma) and cleared by ultracentrifugation at 100,000g at 4 °C for 1 hour in polycarbonate tubes (Beckman Coulter, 343775) in a TLA-100 rotor. The supernatant was sonicated for 10 minutes and then fractionated by medium pH reverse-phase HPLC (Zorbax 300Extend C18, 4.6 x 250 mm column, Agilent) with 10 mM ammonium bicarbonate, pH 8.0, using 5% acetonitrile for 17 minutes followed by an acetonitrile gradient from 5% to 30%. Fractions were collected starting at minute 17 with a flow rate of 0.5 mL/min into a 96 well-plate every 38 seconds. These fractions were pooled into 24 fractions by alternating the wells in the plate as in ^1^.Each fraction was dried and resuspended in 100 μL of HPLC water. Fractions were acidified to pH <2 with HPLC-grade triflouroacetic acid and a stage-tip was performed to desalt the samples.^31^ For LC-MS analysis, samples were resuspended to 1 μg/μL in 1% FA and HPLC-grade water, and ~1 μg of peptides were analyzed per 1 hour run time.

### LC-MS analysis

For HPLC-separation of peptides during MS-analysis, solvent A consisted of 2% DMSO (LC-MS-grade, Life Technologies), 0.125% formic acid (98%+, TCI America) in water (LC-MS-grade, OmniSolv, VWR), solvent B of 80% MeCN (LC-MS-grade, OmniSolv, Millipore Sigma), 2% DMSO and 0.125% formic acid in water.

The following gradients (120 min for unfractionated samples and 90 min for fractionated samples) with percentage of solvent B were applied at a constant flow rate of 350 nL/min: 0% – 12% in 5 min; 12% – 35% in 100 min (unfractionated samples) or 12% – 26% in 70 min (fractionated samples); 35% (or 26%) – 100% in 10 min; 100% for 5 min. For electrospray ionization, 2.6 kV were applied between minutes 1 and 113 (or minutes 1 and 83 for fractionated samples) of the gradient through the column.

### TMTProC analysis

The mass spectrometer was operated to analyze positively charged ions in a data-dependent MS2-mode, recording centroid data with the RF lens level at 60% and the following settings: Full scan: Orbitrap detector, AGC target of 4E5 charges, maximum ion injection time of 50 ms, scan range *m/z* 350-1400 with wide quad. isolation enabled, 120k resolution. Maximum cycle time between Full Scans was set to 3 s.

Following the survey scan, the following filters were applied for triggering MS2-scans: monoisotopic peak selection was enabled (“Peptide Mode”) and additionally, the isolation window was shifted towards the monoisotopic peak *via* a diagnostic routine provided by Graeme McAlister (Thermo Fisher). Isolated masses were excluded for 60 s after triggering with a mass tolerance window of ± 10 ppm, while also excluding isotopes and different charge states of the isolated species. Ions with z = 2 – 3 were analyzed if their *m/z*-ratio was between 500 – 1074 (z = 2+) or 350 – 1381 (z = 3+) to assure visibility of the complementary ion clusters in a normal scan range MS2-scan.

For the MS2-acquisition, the following settings were used: AGC-target was set to 7.5E4 charges and the maximum ion injection times was 80 ms. The quadrupole was utilized for isolation with an isolation width of 0.4 Th, and ions were fragmented with 30% CID-amplitude (10 ms activation time, Activation Q of 0.25). The orbitrap was used for analysis with a normal mass range mode and 30k resolution.

For the acquisition of fractionated samples, the method was adjusted slightly: to trigger a MS2-scan, ions had to show a minimum intensity of 1.9E5 (z = 2) or 4.7E5 (z = 3), the maximum ion injection time was increased to 100 ms and Orbitrap resolution was increased to 50k.

### TMTPro-MS2 analysis

The template provided by Thermo Scientific *via* the method editor for MS2-TMT analysis was adjusted slightly for better comparison with TMTProC.

Full scan: Orbitrap detector, AGC target of 4E5 charges, maximum ion injection time of 50 ms, scan range *m/z* 400-1600 with wide quad isolation enabled, 120k resolution. MS1-spectra were recorded in profile mode. Maximum cycle time between Full Scans was set to 3 s.

Following the survey scan, the following filters were applied for triggering MS2-scans: monoisotopic peak selection was enabled (“Peptide Mode”) and additionally, the isolation window was shifted towards the monoisotopic peak via a diagnostic routine provided by Graeme McAlister (Thermo Fisher). Isolated masses were excluded for 60 s after triggering with a mass tolerance window of ± 10 ppm, while also excluding isotopes and different charge states of the isolated species. Ions with z = 2 – 6 were analyzed if they passed a minimum intensity threshold of 2.5E4 a.u..

For the MS2-acquisition, the following settings were used: AGC-target was set to 1.25E5 charges and the maximum ion injection time was 86 ms. The quadrupole was utilized for isolation with an isolation width of 0.4 Th, and ions were fragmented with 38% HCD-energy. The orbitrap was used for analysis with a fixed first mass of *m/z* 110 and 60k resolution and spectra were recorded in centroid mode.

### TMTPro-MS3 analysis

The template provided by Thermo Scientific *via* the method editor for MS3-SPS-TMT analysis was adjusted slightly for better comparison with TMTProC.

Full scan: Orbitrap detector, AGC target of 4E5 charges, maximum ion injection time of 50 ms, scan range *m/z* 400-1600 with wide quad. isolation enabled, 120k resolution. MS1-spectra were recorded in profile mode. Maximum cycle time between Full Scans was set to 3 s.

Following the survey scan, the following filters were applied for triggering MS2-scans: monoisotopic peak selection was enabled (“Peptide Mode”) and additionally, the isolation window was shifted towards the monoisotopic peak via a diagnostic routine provided by Graeme McAlister (Thermo Fisher). Isolated masses were excluded for 60 s after triggering with a mass tolerance window of ± 10 ppm, while also excluding isotopes and different charge states of the isolated species. Ions with z = 2 – 6 were analyzed if they passed a minimum intensity threshold of 5E3 a.u..

For the MS2-acquisition, the following settings were used: AGC-target was set to 1E4 charges and the maximum ion injection time was 35ms. The quadrupole was utilized for isolation with an isolation width of 0.7 Th, and ions were fragmented with 35% CID-amplitude (10 ms activation time, activation Q of 0.25). The ion trap was used for analysis in the turbo scan rate mode and spectra were recorded in centroid mode.

For triggering MS3-scans for TMT-reporter ion-based quantification, the following filters were applied: Precursor Selection Range: *m/z* 400 – 1600; Precursor Ion Exclusion: *m/z* 50 (low) and *m/z* 5 (high); Isobaric Tag Loss Exclusion (TMTPro). 10 notches were used to isolate SPS-MS3-precursors.

For MS3-scans, the following settings were used: MS1-isolation window: 0.7 Th; MS2-isolation window: 3 Th; ions were fragmented in the HCD cell with a normalized collision energy of 55%. The Orbitrap detector was used, with an AGC target of 1.5E5 charges, maximum ion injection time of 86 ms and 50k resolution. The scan range was set to *m/z* 100 – 500. Spectra were recorded in centroid mode.

### FAIMS analysis

To analyze the influence of field asymmetric ion mobility spectrometry, a FAIMS Pro-device (Thermo Fisher Scientific) was utilized in some experiments. In these experiments, all methods were used as described above, but throughout the experiments, two compensation voltages (−45 V and −65 V) were applied successively to allow the analysis of different ion populations for the specified max. cycle time of 3s. The FAIMS device was operated in standard resolution mode.

### Determination of TMTPro isotopic impurities in balancing and reporter ion parts

To determine the isotopic impurities of each TMTPro tag, we first injected ^13^C_6_ heavy arginine to determine its isotope distribution. This vector is defined as:

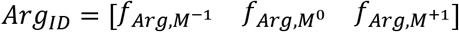

Then, each TMTPro tag was reacted with ^13^C_6_ heavy arginine in a ratio of 5:1 for 2 hours. Labelled heavy arginine was then injected into the mass spec. We define the vector describing the isotope distribution in the MS1 as

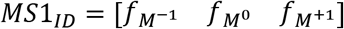

The isotopic impurities of the reporter ion were determined by successively isolating and fragmenting the M^0^, M^+1^, and M^-1^ peaks. The relative signal of each reporter ion in the MS2 completes a matrix for each tag. For example, the 127N tag:

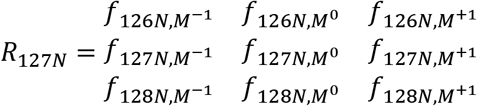

The overall isotopic impurities of the reporter region of this tag are then the dot product of matrix *R_127N_* and the vector *MS*1_*ID*_.

The isotope distribution of the complementary portion of the ions (Heavy Arginine + TMTPro – Reporter) is solved for using the identical method as the reporter ions. The isotope distribution of the heavy arginine is removed by setting up the following system of equations:

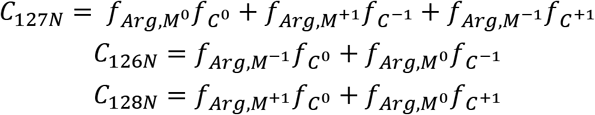

These equations assume that the contribution of isotopes with 2 more or 2 less heavy atoms than the ideal tag is negligible.

Finally, the outer product of the *R*_127*N*_ and [*C*_127*N*_, *C*_128*N*_, *C*_129*N*_]vectors results in a matrix that is used to deconvolve the signals of real peptide distributions (Sup. Fig. S2).

### Data Analysis

The data was analyzed using the Gygi Lab GFY software licensed from Harvard. Thermo .raw-files were converted to mzXML using ReAdW.exe (http://svn.code.sf.net/p/sashimi/code/). Incorrectly assigned precursor charge state as well as incorrectly determined monoisotopic peaks were corrected by custom code.^30^ Assignment of MS2 spectra was performed using the SEQUEST algorithm by searching the data against the combined reference proteomes for Homo Sapiens and S. cerevisiae acquired from Uniprot on 08/07/2016 (SwissProt + Trembl) along with common contaminants such as human keratins and trypsin.

The target-decoy strategy was used to construct a second database of reversed sequences that were used to estimate the false discovery rate on the peptide level. SEQUEST searches were performed using a 20 ppm precursor ion tolerance with the requirement that both N- and C-terminal peptide ends are consistent with the protease specificities of LysC and Trypsin. For high-resolution MS2 data (TMTPro-MS2, TMTProC) the fragment ion tolerance of the MS2 spectrum was set to 0.02 Da, whereas this value was set to 1 Da for low-resolution MS2 spectra acquired with TMTPro-MS3. TMTPro (+ 304.2071 Da) (or, when using TMTPro0, + 295.1896 Da) was set as a static modification on N-termini and lysines residues, and N-ethyl maleimide (+125.047679 Da) was set as a static modification on cysteine residues. Oxidation of methionine (+15.99492 Da) was set as a variable modification, as well as the potential ring-opening of NEM-modified cysteines (+18.010564 Da) when searching the fractionated samples. A peptide level MS2 spectral assignment false discovery rate of 1% was obtained by applying the target decoy strategy with linear discriminant analysis as described previously. Peptides were assigned to proteins and a second filtering step to obtain a 1% FDR on the protein level was applied. Peptides that matched multiple proteins were assigned to the proteins with the most unique peptides.

Peptides identified in TMTPro-MS2 and TMTPro-MS3 experiments were only considered quantified if at least 75% of the signal in the MS1 spectra within the range of the isolation window came from the precursor peak (Isolation Specificity > 0.75). No isolation specificity filters were applied to the TMTProC data. For all TMTProC runs, peptides were filtered if after postsearch calibration, one of the complementary peaks had an error of more than 10 ppm from the median of the other peaks. For all methods, peptides were only considered quantified if the signal to FT noise ratio (S:N) across all channels was greater than 40.

Identification of complementary ion peaks, modelling of the isolation window, and deconvolution of the complementary peaks were performed as previously described.^14^

**Figure S1.**
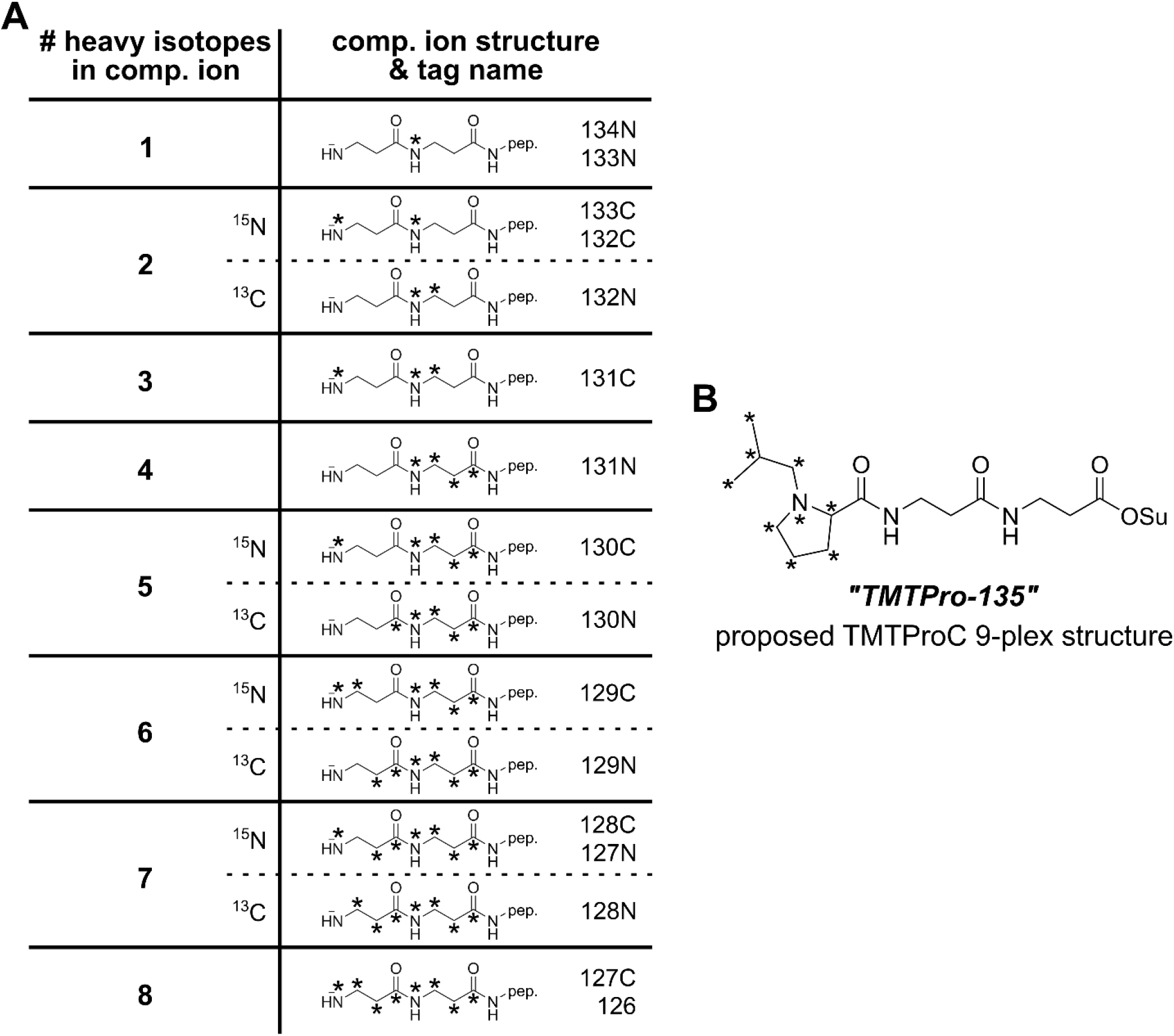
Tag structures and plexing using TMTProC. **A)** TMTProC has up to 8 distinguishable channels using the standard set of 16 tags. Several sets of tags are distinguishable at the low m/z values of the reporter ions because of the difference in increased mass of a heavy carbon and heavy nitrogen atom. **B)** Proposed structure of “TMTPro-135”, which is *quasi-* isobaric to the TMTPro-16plex and extends plexing capacity of TMTPro to nine channels.

**Figure S2.**
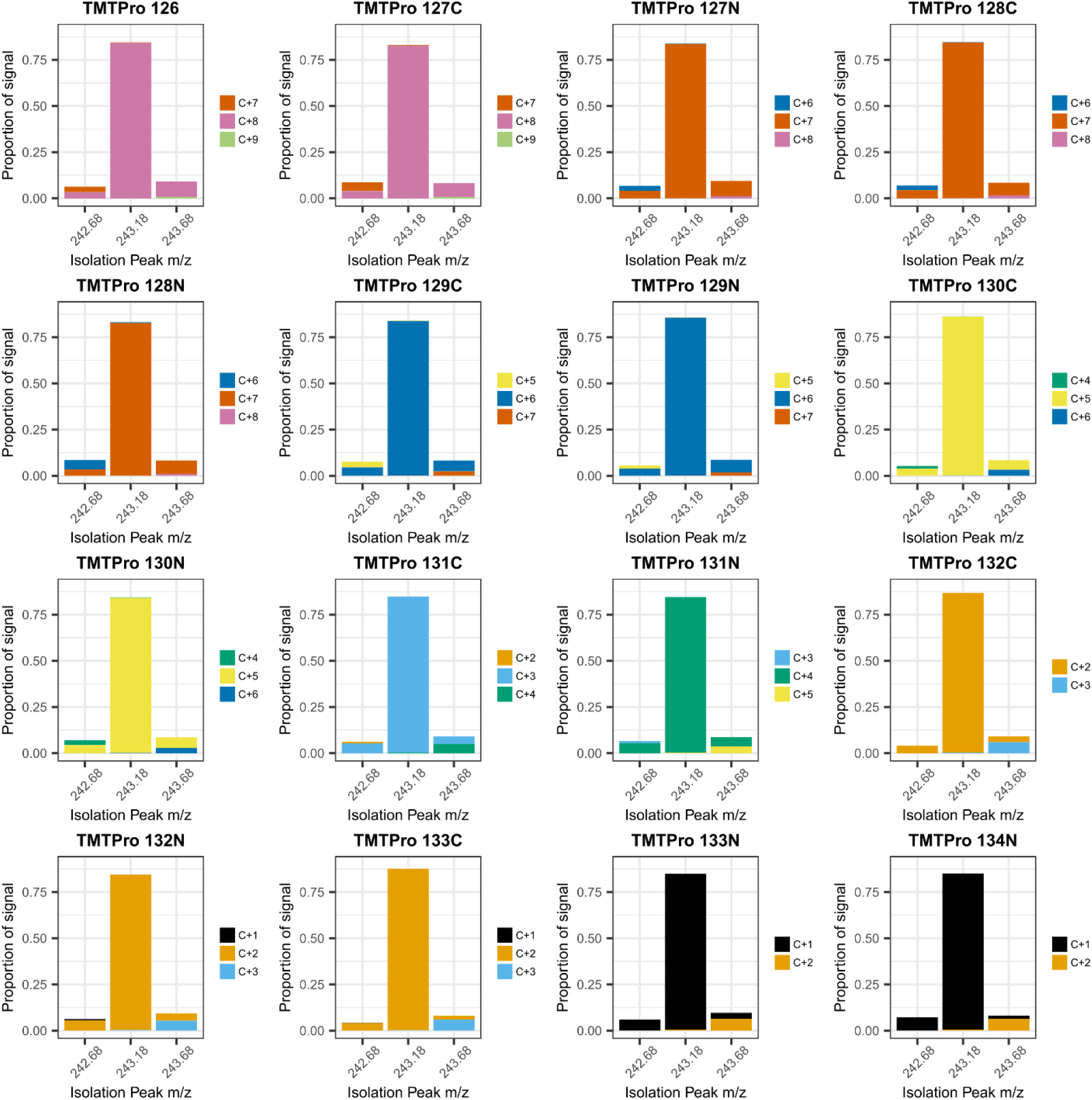
Isotopic impurities for the commercially available TMTPro-tags. To determine the isotopic impurities of each TMTPro tag, we first injected ^13^Cβ heavy arginine to determine its isotope distribution. Then, each TMTPro tag was reacted with ^13^C6 heavy arginine in a ratio of 5:1 for 2 hours. Labelled heavy arginine was then injected into the mass spectrometer. The isotopic impurities of the reporter ion were determined by successively isolating and fragmenting the M^0^, M^+1^, and M^-1^ peaks. The isotope distribution of the heavy arginine is removed and a simple system of 3 linear equations are solved for the isotopic distribution.

**Figure S3.**
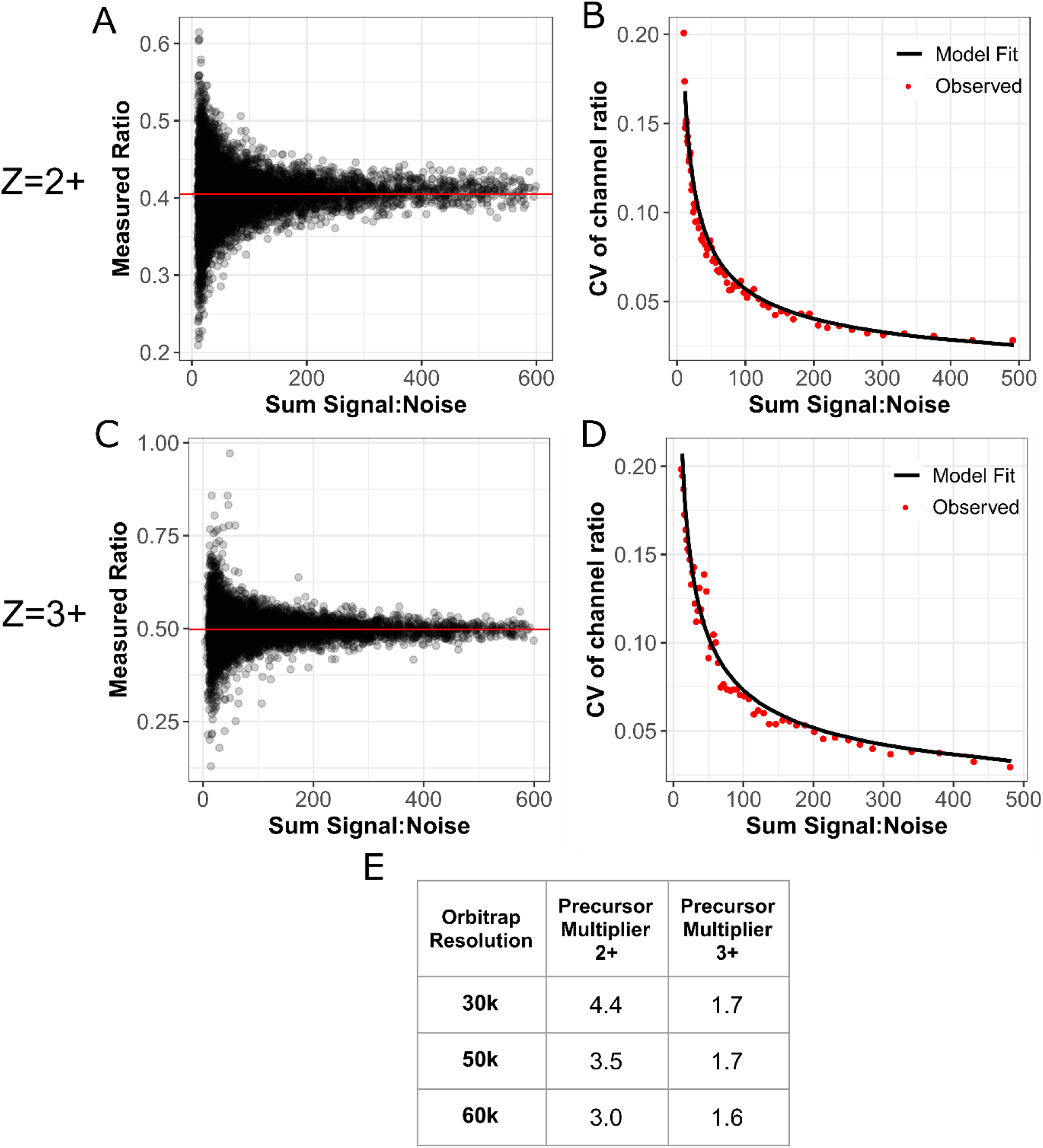
Converting FT signal:noise ratios to pseudo-counts for Bayesian inference. In order to expand the method using Bayesian inference to estimate confidence intervals around protein ratios, we need to determine the conversion factor from signal to FT noise of peptides with a 3+ charge state to pseudo counts. A HeLa sample labelled in ratios of 1:1:1:1:1:1:1:1 with TMTPro was prepared and analyzed either isolating precursors with a 2+ charge state (a-b) or 3+ charge state (c-d). **A,C)** Plots showing how the measured ratio of two channels approaches the true value as the total signal:noise increases **B,D)** The coefficient of variation (CV) of the measured ratio between two channels is plotted as a function of the total Signal:Noise in the two channels. Data are binned and we fit the expected coefficient of variation (CV) of a binomial distribution with a single fitting parameter: the conversion of Signal:Noise to ion counts. **E)** Table of conversion parameters for complementary ions of 2+ and 3+precursors at three different orbitrap resolutions. We find that the conversion of signal:noise into pseudocounts for 3+ peptides is slightly noisier at an Orbitrap resolution of 30k than expected compared to 2+ peptides. The extra noise is likely due to the increased difficulty of deconvolving the complementary ion cluster for 3+ ions.

**Figure S4.**
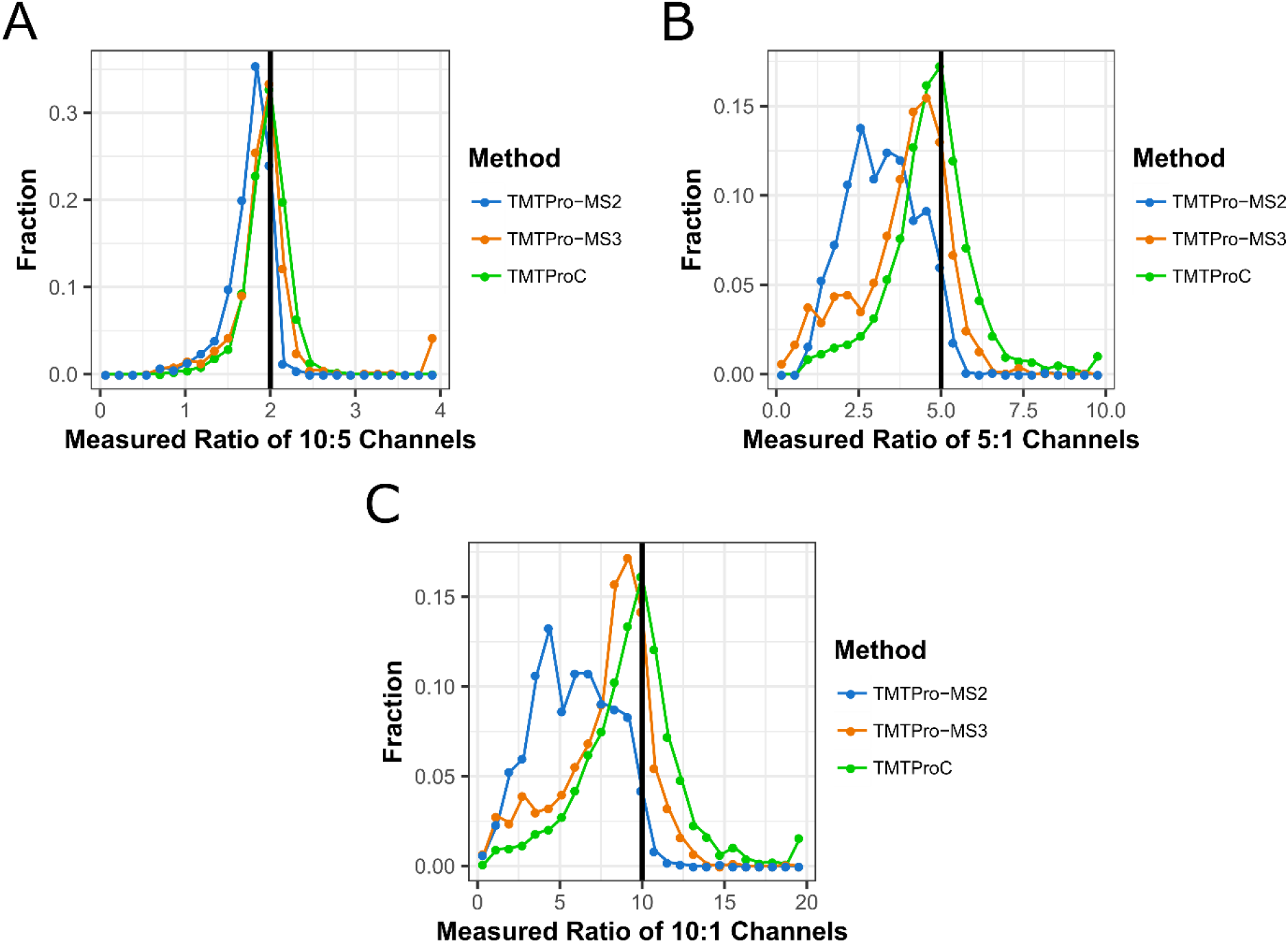
Evaluating ratio distortion for TMTProC and alternative multiplexed quantification strategies with a FAIMS. **A-C)** Experimental design as in Fig. 2. Histograms showing the measured ratios of the 10:5 (a), 5:1 (b), and 10:1 (c) channels for yeast peptides using a FAIMS device (2 CVs) and either TMTPro-MS2, TMTPro-MS3, or TMTProC quantification. Measured ratios more than double their expected value were set to double their expected value. Similar to non-FAIMS runs, MS2 quantification of the 10:1 and 5:1 channels is plagued by interference, which MS3 and TMTProC reduce significantly. In addition, FAIMS improved the mode measured ratio for TMTProC to the actual ratio of the 5:1 and 10:1 channels, while TMPro-MS3 was still subject to some ratio distortion for most peptides with FAIMS.

**Figure S5.**
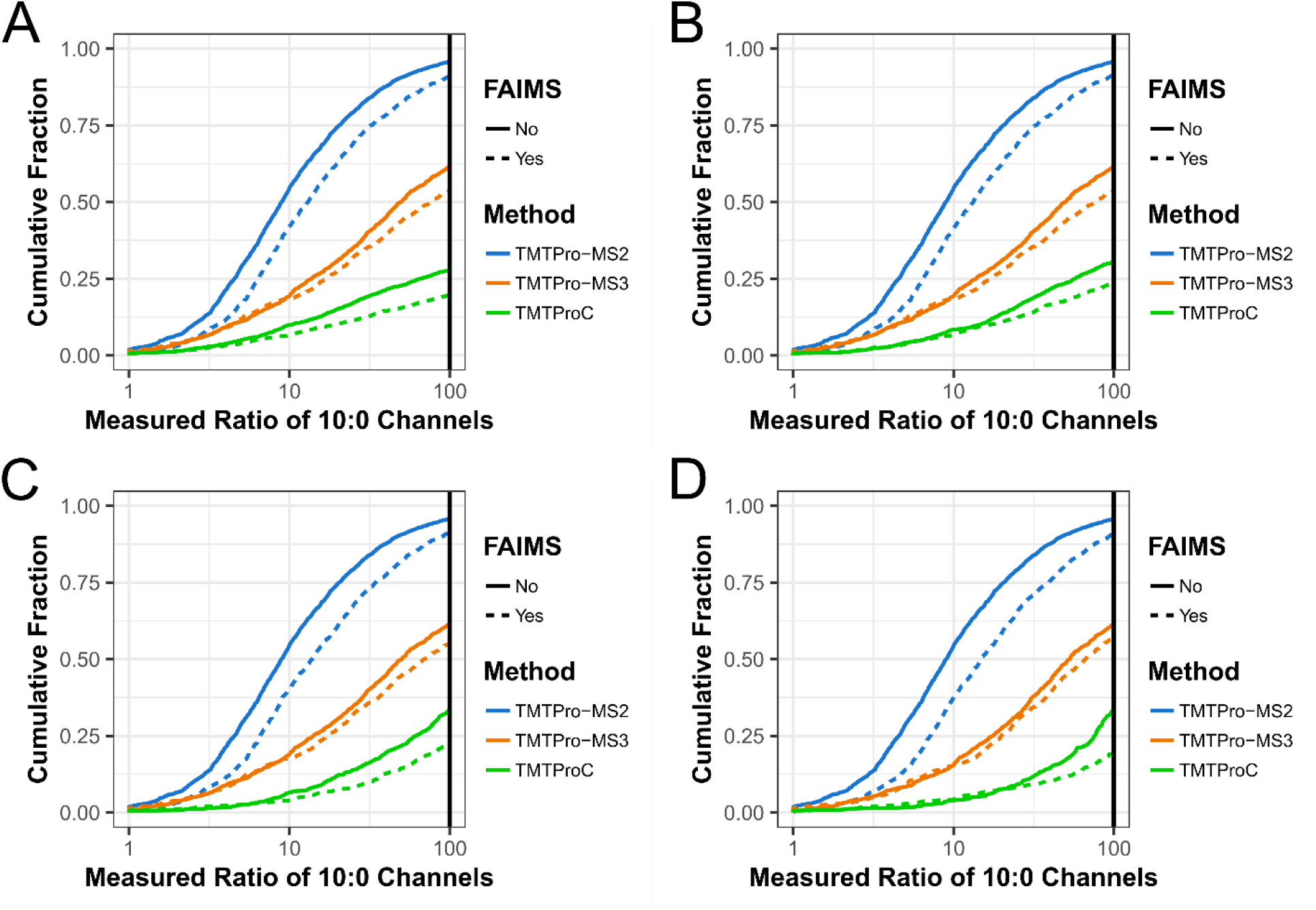
Evaluating ratio distortion for TMTProC and alternative multiplexed quantification strategies with various S:N cutoffs. **A-D)** Experimental design as in Fig. 2. CDF’s are shown for the measured ratio of the 10:0 channels at four different S:N cutoffs: a) 0, b) 200, c) 500 (also shown in Figure 2), and d) 1000. The choice of S:N cutoff does not affect the conclusions of the experiment.

## References

1. Ting L, Rad R, Gygi SP, Haas W. MS3 eliminates ratio distortion in isobaric multiplexed quantitative proteomics. Nat Methods. 2011;8(11):937–940. doi:10.1038/nmeth.1714

2. Thompson A, Wo N, Koncarevic S, et al. TMTpro: Design, Synthesis, and Initial Evaluation of a Proline-Based Isobaric 16-Plex Tandem Mass Tag Reagent Set. Anal Chem. 2019. doi:10.1021/acs.analchem.9b04474

3. Pappireddi N, Martin L, Wühr M. A Review on Quantitative Multiplexed Proteomics. ChemBioChem. 2019;10. doi:10.1002/cbic.201800650

4. Ross PL, Huang YN, Marchese JN, et al. Multiplexed protein quantitation in Saccharomyces cerevisiae using amine-reactive isobaric tagging reagents. Mol Cell Proteomics. 2004;3(12):1154–1169. doi:10.1074/mcp.M400129-MCP200

5. Ong S, Blagoev B, Kratchmarova I, et al. Stable Isotope Labeling by Amino Acids in Cell Culture, SILAC, as a Simple and Accurate Approach to Expression Proteomics *. Mol Cell Proteomics. 2002:376–386. doi:10.1074/mcp.M200025-MCP200

6. Jordan NV, Bardia A, Wittner BS, et al. HER2 expression identifies dynamic functional states within circulating breast cancer cells. Nature. 2016;537. doi:10.1038/nature19328

7. Lignitto L, Leboeuf SE, Homer H, et al. Nrf2 Activation Promotes Lung Cancer Metastasis by Inhibiting the Degradation of Bach1 Article Nrf2 Activation Promotes Lung Cancer Metastasis by Inhibiting the Degradation of Bach1. Cell. 2019;178(2):316–329.e18. doi:10.1016/j.cell.2019.06.003

8. Simsek D, Tiu GC, Flynn RA, et al. The Mammalian Ribo-interactome Reveals Ribosome Functional Diversity and Heterogeneity Article The Mammalian Ribo-interactome Reveals Ribosome Functional Diversity and Heterogeneity. Cell. 2017;169(6):1051–1057.e18. doi:10.1016/j.cell.2017.05.022

9. Ow SY, Salim M, Noirel J, Evans C, Rehman I, Wright PC. iTRAQ Underestimation in Simple and Complex Mixtures: “The Good, the Bad and the Ugly.” J Proteome Res. 2009;8:5347–5355. doi:10.1021/pr900634c

10. Wenger CD, Lee MV, Hebert AS, et al. Gas-phase purification enables accurate, multiplexed proteome quantification with isobaric tagging. Nat Methods. 2011;8(11):1–5. doi:10.1038/nmeth.1716

11. McAlister GC, Nusinow DP, Jedrychowski MP, et al. MultiNotch MS3 enables accurate, sensitive, and multiplexed detection of differential expression across cancer cell line proteomes. Anal Chem. 2014;86(14):7150–7158. doi:10.1021/ac502040v

12. Yu Q, Paulo JA, Naverrete-perea J, et al. Benchmarking the Orbitrap Tribrid Eclipse for Next Generation Multiplexed Proteomics. Anal Chem. 2020. doi:10.1021/acs.analchem.9b05685

13. Wühr M, Haas W, McAlister GC, et al. Accurate multiplexed proteomics at the MS2 level using the complement reporter ion cluster. Anal Chem. 2012;84(21):9214–9221. doi:10.1021/ac301962s

14. Sonnett M, Yeung E, Wühr M. Accurate, Sensitive, and Precise Multiplexed Proteomics using the Complement Reporter Ion Cluster. Anal Chem. 2018.

15. Hart EM, Gupta M, Wühr M, Silhavy TJ. The gain-of-function allele bamA E470K bypasses the essential requirement for BamD in β-barrel outer membrane protein assembly. Proc Natl Acad Sci. 2020;117(31). doi:10.1073/pnas.2007696117

16. Cao WX, Kabelitz S, Gupta M, et al. Article Precise Temporal Regulation of Post-transcriptional Repressors Is Required for an Orderly Drosophila Maternal-to-Zygotic Transition ll Precise Temporal Regulation of Post-transcriptional Repressors Is Required for an Orderly Drosophila Maternal-to. Cell Rep. 2020;31. doi:10.1016/j.celrep.2020.107783

17. Li A, Mao D, Yoshimura A, et al. Multi-Omic Analyses Provide Links between Low-Dose Antibiotic Treatment and Induction of Secondary Metabolism in Burkholderia thailandensis. Mol Biol Physiol. 2020;11(1).

18. Hart EM, Gupta M, Wühr M, Silhavy TJ. The synthetic phenotype of Δbamb Δbame double mutants results from a lethal jamming of the bam complex by the lipoprotein RcsF. Mol Biol Physiol. 2019;10(3):1–12. doi:10.1128/mBio.00662-19

19. Stadlmeier M, Bogena J, Wallner M, Martin W, Carell T. A Sulfoxide-Based Isobaric Labelling Reagent for Accurate Quantitative Mass Spectrometry Angewandte. Angew Chemie. 2018;08544:2958–2962. doi:10.1002/anie.201708867

20. Winter SV, Meier F, Wichmann C, Cox J, Mann M, Meissner F. EASI-tag enables accurate multiplexed and quantification. Nat Methods. 2018;15(July). doi:10.1038/s41592-018-0037-8

21. Peshkin L, Gupta M, Ryazanova L, Wühr M. Bayesian Confidence Intervals for Multiplexed Proteomics Integrate Ion-statistics with Peptide Quantification Concordance. Mol Cell Proteomics. 2019;18(10):2108–2120. doi:10.1074/mcp.TIR119.001317

22. Gupta M, Sonnett M, Ryazanova L, Presler M, Wühr M. Quantitative Proteomics of Xenopus Embryos I, Sample Preparation. In: Xenopus. ; 2018:175–194. doi:https://doi.org/10.1007/978-1-4939-8784-9_13

23. Wessel D, Flugge UI. A Method for the Quantitative Recovery of Protein in Dilute Solution in the Presence of Detergents and Lipids. Anal Biochem. 1984;143:141–143.

24. Hebert AS, Prasad S, Belford MW, et al. Comprehensive Single-Shot Proteomics with FAIMS on a Hybrid Orbitrap Mass Spectrometer. Anal Chem. 2018;90. doi:10.1021/acs.analchem.8b02233

25. Perez-Riverol Y, Csordas A, Bai J, et al. The PRIDE database and related tools and resources in 2019: improving support for quantification data. Nucleic Acids Res. 2019;47:442–450. doi:10.1093/nar/gky1106

26. Kelstrup CD, Aizikov K, Batth TS, et al. Limits for Resolving Isobaric Tandem Mass Tag Reporter Ions Using Phase-Constrained Spectrum Deconvolution. J Proteome Res. 2018; 17. doi:10.1021/acs.jproteome.8b00381

27. Kozhinov AN, Tsybin YO. Filter diagonalization method-based mass spectrometry for molecular and macromolecular structure analysis. Anal Chem. 2012;84(6):2850–2856. doi:10.1021/ac203391z

28. Olsen J V, Macek B, Lange O, Makarov A, Horning S, Mann M. Higher-energy C-trap dissociation for peptide modification analysis. Nat Methods. 2007;4(9):709–712. doi:10.1038/NMETH1060

29. Schwartz JC, Senko MW. A Two-Dimensional Quadrupole Ion Trap Mass Spectrometer. Am Soc Mass Spectrom. 2002;13:659–669.

30. Schweppe DK, Eng JK, Yu Q, et al. Full-Featured, Real-Time Database Searching Platform Enables Fast and Accurate Multiplexed Quantitative Proteomics. J Proteome Res. 2020;19. doi:10.1021/acs.jproteome.9b00860

31. Rappsilber J, Mann M, Ishihama Y. Protocol for micro-purification, enrichment, pre-fractionation and storage of peptides for proteomics using StageTips. Nat Protoc. 2007;2(8). doi:10.1038/nprot.2007.261

